# Single cell transcriptomic profiling identifies tumor-acquired and therapy-resistant cell states in pediatric rhabdomyosarcoma

**DOI:** 10.1101/2023.10.13.562224

**Authors:** Sara G Danielli, Yun Wei, Michael A Dyer, Elizabeth Stewart, Marco Wachtel, Beat W Schäfer, Anand G Patel, David M Langenau

## Abstract

Rhabdomyosarcoma (RMS) is a pediatric tumor that resembles undifferentiated muscle cells; yet the extent to which cell state heterogeneity and molecular features are shared with human development have not been fully ascribed. Here, we report a single-cell/nucleus RNA sequencing atlas derived from 72 datasets that includes patient tumors, patient-derived xenografts, primary *in vitro* cultures, and established cell lines. We report four dominant muscle-lineage cell states in RMS: progenitors, proliferative, differentiated, and ground cells. We stratify these RMS cells along the continuum of human muscle development and show that RMS cells share expression patterns with fetal/embryonal myogenic precursors rather than postnatal satellite cells. Indeed, fusion-negative RMS (FN-RMS) have a discrete stem cell hierarchy that faithfully recapitulates fetal muscle development. We also identify therapy-resistant FN-RMS progenitor cells that share transcriptomic similarity with bipotent skeletal mesenchymal cells, while a subset of fusion-positive (FP) RMS have tumor-acquired cells states, including a neuronal cell state, that are not found in development. Chemotherapy induced upregulation of progenitor signatures in FN-RMS while the neuronal gene programs were retained after therapy in FP-RMS. Taken together, this work identifies new cell state heterogeneity including unique treatment-resistant and tumor-acquired cell states that differ across RMS subtypes.

## INTRODUCTION

Rhabdomyosarcoma (RMS) is a pediatric solid tumor that shares features with arrested skeletal muscle precursors [1, 2]. Pediatric RMS has been classified into three major subtypes that have divergent molecular drivers: (i.) fusion-positive RMS (FP-RMS) have DNA translocations that juxtapose *PAX3* or *PAX7* with *FOXO1* (PAX3::FOXO1 or PAX7::FOXO1); (ii.) fusion-negative RMS (FN-RMS) lack pathognomonic translocation events, but often have oncogenic activation of RAS signaling; and (iii.) spindle cell/sclerosing rhabdomyosarcoma (SS-RMS, a subclass previously classified as FN-RMS) are largely driven by NCOA2::VGLL2 translocations or a p.Leu122Arg mutation in the *MYOD1* transcription factor [3–13]. Despite aggressive therapies combining radiation, chemotherapy, and surgery, 70% of patients with unresectable or disseminated disease develop recurrent RMS that has a dismal 5-year overall survival rate of under 20% [14–17].

Like many pediatric cancers, RMS tumors have a low mutational burden and few known genetic alterations reliably predict recurrent disease [8, 18–20]. Thus, it is critical to understand the non-genetic heterogeneity within RMS, and the role that specific cell subpopulations play in driving the clinical behavior of RMS. Indeed, multiple groups have applied single-cell transcriptomics to discover distinct RMS cell subpopulations [21–24]. These studies consistently identified malignant cells with expression patterns similar to developing skeletal muscle, yet each study introduced different nomenclature and classification strategies due to limited sample numbers, differences in bioinformatic approaches, and mapping shared developmental cell states across mouse and/or human muscle. As a result, there is a need to clearly define cell states, to assess developmental similarity between RMS and human muscle, and to evaluate the dynamics of cell state transitions during therapy.

Here, we present a consensus evaluation of intratumoral heterogeneity in human RMS by combining datasets encompassing 72 samples from patient tumors, patient-derived xenograft (PDX) models, PDX-derived primary cell cultures (PDCs), and commercial cell lines (CLs) [21–23, 25]. By uniformly processing and integrating these datasets, we generated a comprehensive and unified annotation of RMS-specific cell subpopulations. In total, we identified four major RMS cell subpopulations – (1) progenitor cells that are largely quiescent and express characteristic mesenchymal and extracellular matrix genes; (2) differentiated cells that are post-mitotic and resemble mature skeletal muscle; (3) proliferative cells that are actively dividing but largely lack expression of progenitor or differentiated cell programs; and (4) ground state cells that lack expression of the other three dominant signatures. While we identified shared RMS cell states with embryonal and fetal skeletal muscle development, we also found subtype-specific cell states. FP-RMS include a unique neuronal cell state, indicating that FP-RMS acquire non-myogenic gene expression programs during tumorigenesis. In addition, progenitor cells in FN-RMS closely resemble bipotent SkM.Mesenchymal cells found in fetal muscle development, which was not observed in FP-RMS. Both FN-RMS and FP-RMS failed to share similarity with postnatal satellite cells. Together, these results challenge the dogma that RMS follow rigid muscle developmental hierarchies and RMS originate from or resemble satellite-cell derived post-natal muscle. Finally, we show that our cell state signatures can be used to identify treatment-persistent cell populations. Specifically, progenitor and neuronal signatures were significantly enriched in treated samples in FN-RMS and FP-RMS, respectively. In total, this work presents a harmonized model of intratumoral heterogeneity within RMS and provides new insights into the intersection of normal development and therapy resistance within cancer.

## RESULTS

### A single-cell/nucleus transcriptomic atlas of RMS

Several groups have investigated the transcriptional heterogeneity of rhabdomyosarcoma (RMS) using single-cell RNA (scRNAseq) and/or single-nucleus RNA sequencing (snRNAseq). These studies identified RMS cell states using different bioinformatic methods, leading to divergent and often confusing nomenclature [21–23]. To overcome these limitations, we collected and uniformly processed 72 scRNAseq or snRNAseq datasets from four previously published studies that used the 10X Genomics platform [21–23, 25] (*n* = 107,523 malignant cells/nuclei). This unified cohort included tumors and experimental models derived from patients seen across four medical centers worldwide, along with established cell line models. This dataset encompasses the largest transcriptomic atlas for any sarcoma analyzed to date and includes patient tumors (*n* = 21), PDXs (patient-derived xenograft, *n* = 32), PDCs (patient derived cell culture, *n* = 14), and conventional CLs (*n* = 5) (**Figure 1A**). Forty-five datasets were generated from either patient tumor samples or PDXs from the St. Jude Childhood Solid Tumor Network (CSTN)[26], and an additional 6 PDCs were generated from CSTN xenografts (Table S1)[23]. Importantly, these samples are representative of intermediate and high-risk RMS, and include primary, recurrent, and metastatic tumors (**Table S1**). All major subtypes of disease were represented including FP-RMS (*n* = 27, previously known as alveolar RMS), FN-RMS (*n* = 43, previously known as embryonal RMS), and two SS-RMS cases with *MYOD1*^L122R^-mutations (**Table S1**). After merging datasets and performing dimensionality reduction, malignant cells/nuclei grouped separately based on patient and model systems, consistent with observations in other cancer types [27–29] (**Figure 1B** **and S1A**).

**Fig. 1.**
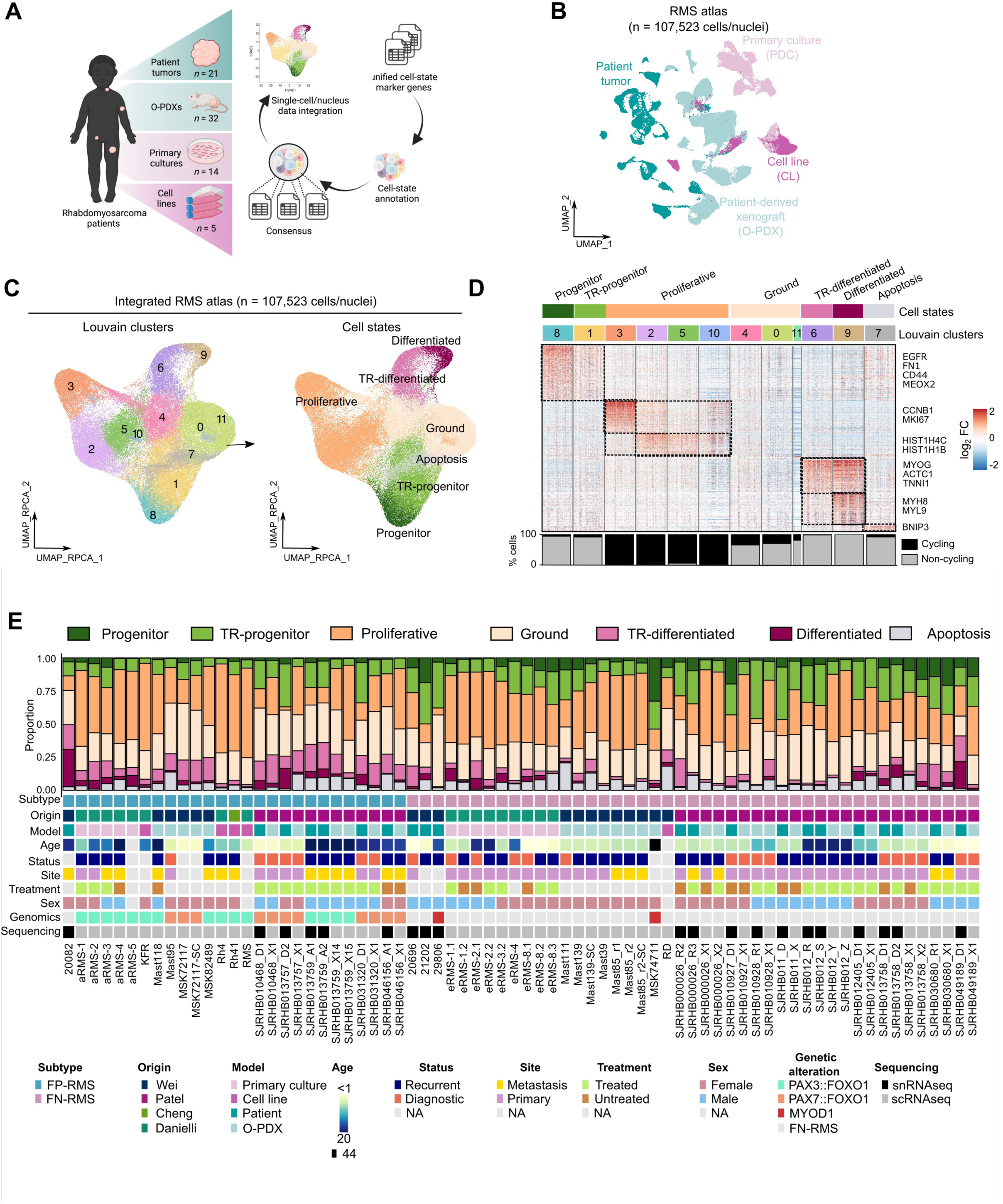
Integrated analysis of single cell and single nuclei sequencing identifies dominant cell states in human RMS. **A**) Schematic of approach and RMS models profiled by single-cell analysis. **B)** UMAP plot of tumor cells/nuclei (n = 72 datasets) colored by model of origin [21–23]. **C)** UMAP plot of integrated tumor cells/nuclei using reciprocal PCA (RPCA) projection. Cells/nuclei are colored by Louvain cluster (left) or assigned cell states (right). **D)** Heatmap of the genes (x axis) enriched in Louvain clusters (y axis) across the integrated RMS dataset (FC>0.25; *n* = 400 representative cells/nuclei shown with exception of cluster 11 that contained *n* = 101 cells/nuclei). The percentage of cycling cells within each cluster shown in the bar plot below. **E)** Summary of the cell state composition of each RMS dataset (n = 72) with clinical information included as an oncoplot below. FN-RMS, fusion-negative RMS; FP-RMS, fusion-positive RMS; PDC, primary culture; CL, cell line; PDX, patient-derived xenograft; snRNAseq, single-nuclei RNA sequencing; scRNAseq, single-cell RNA sequencing; NA, not available; TR-progenitor, transiting-progenitor; TR-differentiated, transiting-differentiated. Gene lists used for generating panels D and E are shown in Table S2.

To identify shared transcriptomic signatures across different samples, we corrected for inter-patient variation utilizing anchor-based integration [30]. Following integration, samples were intermixed and as expected, we did not identify outlier cells/nuclei that were attributable to only one patient, dataset, or model system (**Figure S1B**). We next applied unsupervised clustering and identified 12 Louvain clusters that could be grouped into distinct subpopulations based on shared transcriptomic profiles (**Figure 1C-D; Figure S1C**). These subpopulations include (1) two clusters with cells/nuclei expressing varying levels of mesoderm transcription factors (e.g. *MEOX2*), cell surface markers (e.g. *CD44*, *EGFR*), and extracellular matrix proteins (e.g. *FN1*) which we call the “progenitor” and “transiting-progenitor” (“TR-Progenitor”) that were distinguished from each other based on overall levels of marker expression; (2) a “proliferative” subpopulation that comprised four clusters that shared GSEA signature similarity with proliferative and DNA replication gene modules; (3) two clusters of “transiting-differentiated” (“TR-Differentiated”) and “differentiated” muscle cells/nuclei expressing transcription factors from committed muscle cells (e.g. *MYOG*) and muscle contraction proteins (e.g. *TNNI1*, *MYH8*); (4) an “apoptotic” subpopulation expressing genes associated with cell death (e.g. *BNIP3*); and (5) “ground” cells that do not show any enrichment of these signatures (**Figure 1C-E** **and S1C**; **Table S2 and 3**). Importantly, we compared five matched PDX samples that were generated by the St. Jude Childhood Solid Tumor Network [26] and that were independently expanded and processed in different labs [21, 22]. We noted consistency in cell subpopulation distributions for these samples derived from the same patient, indicating that our analysis was not skewed by experimental setting, xenograft passaging, or protocol differences in cell isolation for scRNAseq (**Figure S1D)**. One important new discovery of this analysis was the existence of transitioning cell states that had overall lower levels of transcriptionally defined cell states genes (‘TR-progenitor’ and ‘TR-differentiated’ cells), suggesting that RMS cells exist along a continuum of gene expression from less differentiated progenitor cells to differentiated muscle-like cells.

### A tripartite cell state landscape of RMS

In previous reports, each group identified clusters of cells/nuclei with transcriptomic similarity to muscle-lineage cells [21–23]. Despite these similarities, each used different sample types, computational methods to define gene expression signatures, and differing nomenclature for each subpopulation. For example, Patel *et al.* analyzed 18 matched samples from primary patient and orthotopic PDX models and identified three cell populations which they called mesoderm, myoblast, and myocyte cells based on perceived similarity with mouse muscle development [21]. In their study, the myoblast compartment included both proliferative and non-proliferative cells. In contrast, Wei et al. examined 9 PDX and 4 patient tumors to identify four RMS-specific cell subpopulations that they called mesenchymal, proliferative, differentiated, and ground cells [22]. Finally, Danielli *et al.* studied 14 PDCs and 3 conventional cell lines to group RMS cells into muscle stem-cell-like cells, cycling progenitors, and differentiated cells [23].

We leveraged our unified RMS cell atlas to refine these cell state signatures. We scored the single-cell and single-nucleus profiles within the integrated RMS atlas using published signatures from each prior study. Signatures from all three studies identified similar patterns of heterogeneity in the progenitor-like and differentiated-like cells (**Figure 2A**). The one exception was the myoblast signature from Patel et al. which was broadly expressed in most RMS cells or nuclei within the unified atlas. Collectively, we detected significant overlap between gene signatures identified across the three datasets indicating that the previously published studies had independently uncovered largely similar dominant RMS cell states. These data are also consistent with recently published findings from DeMartino et al. that defined a tripartite cell state landscape in FN– and FP-RMS using a different nomenclature [24].

**Fig. 2.**
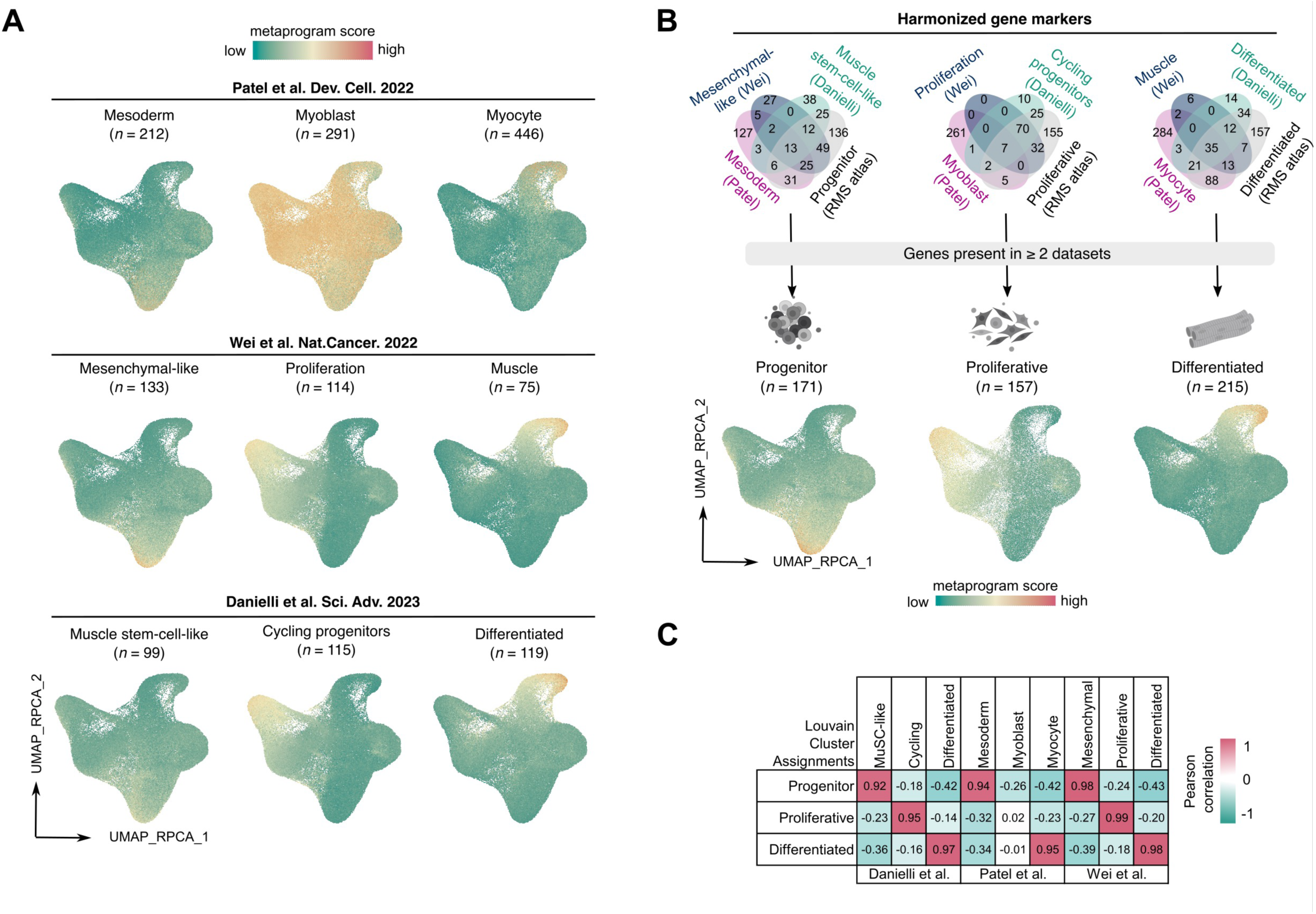
A tripartite cell state landscape of RMS cells and identification of cell state RMS metaprograms. **A**) UMAP plots of integrated tumor cells/nuclei (*n* = 107,523 cells) scored for the metaprograms identified in the original publications. Number of genes within each metaprogram noted. **B)** Comparison of published cell state metaprograms and those defined by our Louvain clustering approach. Top: Venn diagrams showing overlap of gene markers across the three original publications and our new analysis (“RMS atlas”). Bottom: UMAP plots of integrated tumor cells/nuclei (*n* = 107,523 cells) showing expression of the newly defined, high confidence cell state gene signatures. Number of genes within each metaprogram noted. **C)** Pearson correlation coefficients for the metaprograms identified in the three original publications and the new metaprogram signatures defined by our work.

To construct consensus signatures for the three dominant RMS cell states, we selected genes that were present in at least two datasets to generate signatures for each of the major RMS cell subpopulations. We generated three signatures that we call progenitor (*n* = 171 genes), proliferative (*n* = 157 genes), and differentiated (*n* = 215 genes) signatures (**Figure 2B; Table S4)**. Because these unified gene signature lists were generated from a variety of models (patient tumors, PDXs, PDCs and CLs), they represent a robust and broadly applicable set of markers for defining each RMS cell state. As expected, these new high-confidence consensus cell state signatures demonstrated significant overlap with those originally reported, with the exception of the Patel, *et al.* myoblast signature (**Figure 2C**).

### The muscle lineage score stratifies between RMS subtypes

Unsupervised clustering identified previously unknown transitory cells within RMS (**Figure 1C-E**), which led us to test whether tumor cell heterogeneity exists within a continuum between progenitor and differentiated cell states. We created a “muscle lineage score,” defined as the difference between the differentiated and progenitor signature scores and applied this scoring metric to every single-cell/nucleus profile within our atlas and mapped this in relation to our proliferation score. Indeed, there was significant heterogeneity within tumors based on stratifying cells using the muscle lineage score. Yet, unexpectedly, FP-RMS samples had an overall higher muscle-lineage score when compared to datasets derived from FN-RMS, both at the single-cell and pseudo-bulk level (**Figure 3A-C**). Also of note, FP-(n=93) or FN-(n=67) core signatures reported by Wei et al.[22] also largely stratified these two subtypes (**Figure S2B**). Together, these findings suggest that FP-and FN-RMS, despite sharing largely similar cell states, also have activation of subtype specific gene programs that are found within all tumor cells.

**Fig. 3.**
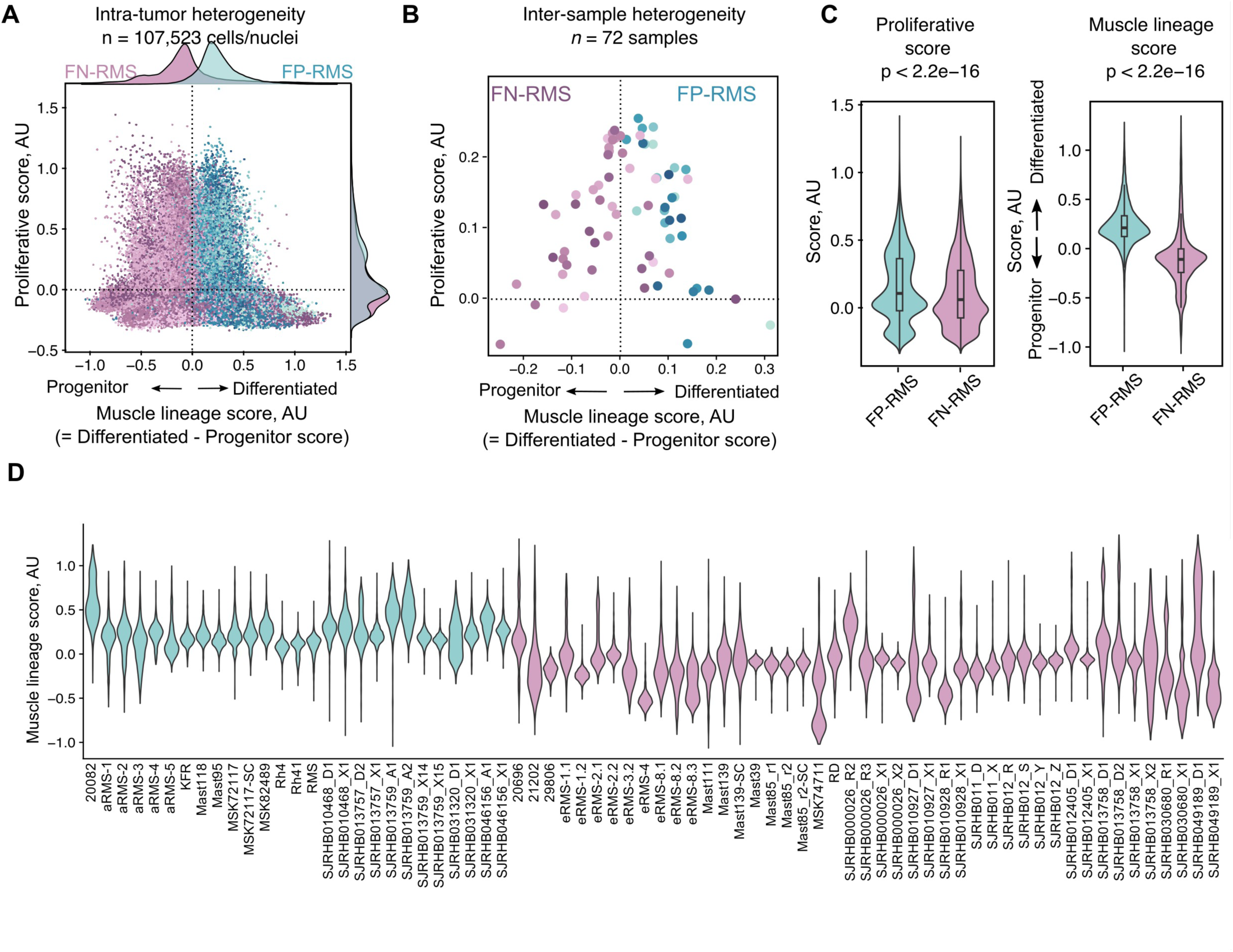
RMS cells lie in a continuum of gene expression defined by three dominant cell states including progenitor, differentiated, and proliferative cell states. **A**) Graphical analysis showing expression of all RMS cells based “muscle lineage score,” defined as the difference between the differentiated and progenitor signature scores and proliferation score. Subtypes denoted by purple (FN-RMS) and blue (FP-RMS). **B)** Average cell expression based on pseudo-bulk analysis for each of the 72 RMS samples. **C)** Violin plots showing individual cell expression of proliferative (left) and muscle lineage score (right) across FP-RMS (*n* = 40,526) and FN-RMS (*n* = 69,997) cells. **D)** Violin plots showing cell expression of the muscle-lineage score across each of the 72 RMS samples. FN-RMS, fusion-negative RMS; FP-RMS, fusion-positive RMS; AU, arbitrary unit. Statistics used Student’s t-test with p-values noted in the figure.

Most cycling cells/nuclei had an intermediate muscle lineage score that was irrespective of subtype, suggesting that they largely lacked expression of either progenitor or differentiated genes (**Figure 3A-C** **and S2A)**. We also observed considerable intertumoral variability, particularly in FN-RMS samples (**Figure 3D**), where we identified samples with exceptionally low lineage scores (e.g., MSK74711, a *MYOD1^L122R^*-mutant FN-RMS sample, and SJRHB010928_R1, a pre-treated FN-RMS tumor sample) or samples with high lineage scores (e.g., SJRHB00026_R2 and SJRHB049189_D1). In total, our data support a model where RMS cells lie in a continuum of gene expression defined by three dominant cell states including progenitor, proliferative, and differentiated cell states while also containing subtype specific gene programs found in all RMS cells within the tumor.

### Neuronal cells are a unique feature of FP-RMS

Our initial analyses centered on combining RMS subtypes together to identify conserved cell states shared across pediatric RMS. While this approach enabled us to define key muscle-lineage cell states shared across RMS, it would likely fail to identify subtype-specific subpopulations within the RMS atlas. Our combined large cohort of RMS samples also enabled us to evaluate heterogeneity within PAX3::FOXO1 and PAX7::FOXO1 translocated tumors as distinct entities. As expected, we identified progenitor, proliferative and differentiated cell subpopulations in each molecular subtype (**Figure 4A** and **S3A**). Yet, when we analyzed the overlap in expressed genes from these seemingly shared states, we observed that progenitor cell states were not transcriptionally the same across each tumor subtype (**Figure S3B**). In addition, we also found unique subtype-specific clusters, which we annotated using gene set enrichment analysis and cluster-specific gene expression (**Table S5-6**). Namely, we found: (1) a group of cells/nuclei in FN-RMS that express interferon response genes such as *ISG15* and *IFIT1-3* (“IFN” cluster; 1.5% of total cells/nuclei), and (2) a small FP-RMS cluster that express neuronal genes including *SOX11*, *DCX*, *L1CAM*, and *CHGA* (‘neuronal’ cluster; 1.4% and 4.8% of total cells/nuclei from PAX3::FOXO1 and PAX7::FOXO1 FP-RMS, respectively, **Figure 4A**). Importantly, cells/nuclei expressing the neuronal gene signature were detected in only FP-RMS samples (*n* = 5 of 11 PAX7::FOXO1 FP-RMS and *n* = 5 of 15 PAX3::FOXO1 FP-RMS, defined as > 1% of total cells/nuclei; **Table S7**). Gene set enrichment analysis confirmed that the neuronal cluster was enriched for genes associated with neurogenesis pathways including axonogenesis (GO:0007409), central nervous system development (GO:0007417) and central nervous system neuron differentiation (GO:0021953; **Table S6**). Despite tumors retaining the tripartite muscle lineage programs across models (**Figure S3C**), neuronal subpopulations were detected in larger numbers in patient tumors and PDXs, but rarely or not at all in primary cultures or commercial cell lines [**Figure S4; Table S7**]. The absence of neuronal cells within cell lines may explain why this rare subpopulation was not identified previously by scRNA sequencing analysis by Danielli *et al.*[23], and may indicate that PDXs are the appropriate experimental model for better clarifying the function of neuronal cells in FP-RMS. Patel et al. and DeMartino et al. also failed to identify these neuronal-pathway enriched tumor cell states, likely reflecting the use of bioinformatic approaches that combined subtypes together for defining cell states [21, 24].

**Fig. 4.**
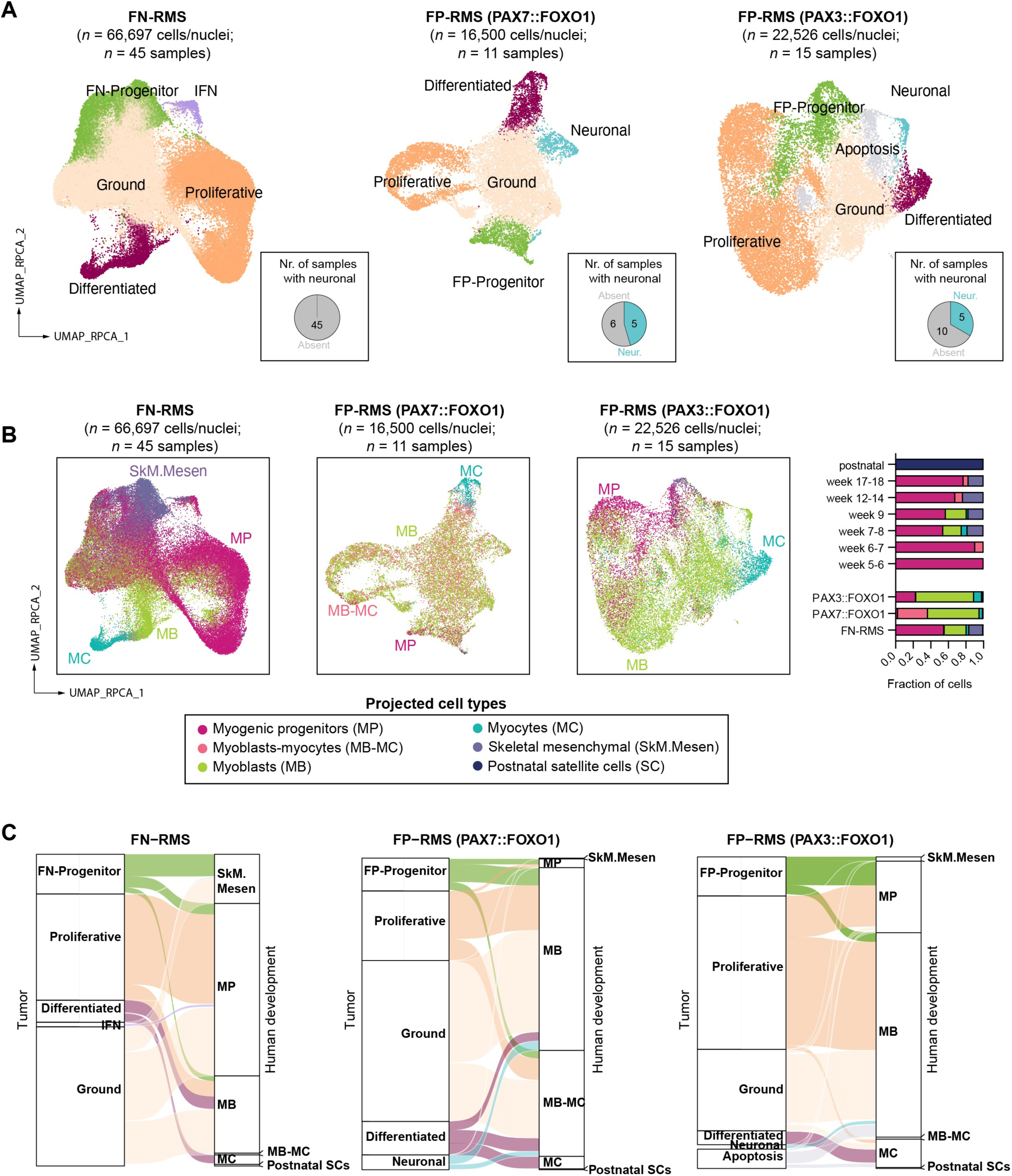
Subtype analysis reveals shared RMS cell heterogeneity with human skeletal muscle development and novel tumor-derived cells states. **A**) UMAP plots of FN-RMS, PAX7::FOXO1 and PAX3::FOXO1 FP-RMS. Cells/nuclei were integrated independently and colored based on cell state. The number of samples having ≥1% neuronal cells are shown in the pie charts at the bottom right of each graph. **B)** Comparison of RMS cell state heterogeneity with muscle cell types found in human skeletal muscle development as defined by Xi *et al.* 2020 (*Cell Stem Cell*). Left: UMAP plots for each subtype. Right: Quantification of cell types found within developmentally defined ages (top) and RMS subtypes (bottom). **C)** Sankey plots showing the proportion of tumor cells classified according to their most similar human developmental equivalent.

### Mapping shared cell heterogeneity between RMS and human skeletal muscle development

Human skeletal myogenesis proceeds in three defined waves [31]. First, mesodermal progenitors (MPs) from the somite create embryonic myoblasts (MBs) and myocytes (MCs) that drive early skeletal muscle deposition. A second wave of muscle development occurs coincident with the transition of embryonic to fetal development, where both MP and bipotent skeletal muscle mesenchymal progenitor (SkM.Mesen) cells likely drive muscle formation [31, 32]. Finally, a third wave of muscle growth is regulated by classically defined muscle stem cells called satellite cells that expand during early postnatal growth to create muscle and then become a reserve stem cell population later in life to aid in repair after injury [33–35]. Here, we took advantage of cell annotations from normal human myogenic development [31] to compare RMS cells/nuclei to these established myogenic cell states (**Figure S5A**).

We used transfer learning with SingleR [36], a computational framework that takes a dataset with known labels as an input and then transfers them onto a test dataset based on similarity to the reference. We show that FN-RMS tumor cells shared similarity with a variety of developing human muscle cell types including MP, SkM.Mesen, MB, and MC (**Figure 4B-C; Figure S5B; Table S8**). FN-RMS have shared cell states with those found in the second wave of muscle development that starts at week 7-8. By contrast, most PAX3::FOXO1 and PAX7::FOXO1 FP-RMS shared cell state similarity with MBs and/or myoblasts-myocytes (MB-MCs), with only a minority of cells mapping to MCs or MPs, and few to no cells sharing similarity with SkM.Mesen cells (**Figure 4B-C; Table S8**). FP-RMS cells did not map to a distinct wave of muscle development, raising the possibility that FP-RMS do not strictly adhere to the shared developmental hierarchies found in normal muscle development. Rather, they appear to stochastically adopt muscle cell states across embryonic and fetal development and can also have unique tumor acquired cell states including neural pathway enriched states (**Figure 4**). Based on these findings, we propose a refined nomenclature based on shared developmental similarity (or not) with human muscle development: “FN-skeletal muscle mesenchymal-like” (FN-SkM.Mes-like) for FN-RMS that are similar to bi-potent stem cells found in fetal development; and “FP-progenitor” and “FP-neural” for cells that represent tumor-acquired cell states in FP-RMS. Finally, RMS do not contain appreciable numbers of cells with shared similarity to postnatal satellite cells and do not map to the third wave of muscle development (**Figure 4B-C; Table S8**).

### Identification of therapy-resistant cells states in RMS

Treatment recurrence is a major hurdle to achieving durable long-term treatment responses in RMS, and we reasoned that RMS cell states might also define therapy persistent tumor cells. We evaluated tissue from patients who underwent delayed resection, for whom both pre-treatment and on-treatment samples were available [21]. These unique samples allowed us to study therapy-induced transcriptomic shifts. We generated RNA-sequencing libraries from those samples and applied RMS signatures for the three dominant cell states (**Figure 5A**, *n* = 9, [*n* = 7 FN-RMS, *n* = 2 FP-RMS]). We uncovered a significant increase in progenitor scores in the recurrent/relapsed patient tumors as compared to samples from primary diagnostic biopsy (p=0.02; Wilcoxon signed rank test) and a decrease in proliferative scores (p=0.02; Wilcoxon signed rank test; **Figure 5B**). We also analyzed snRNAseq obtained from a FN-RMS patient before and during therapy, SJRHB00026_R2 and _R3, and noted a significant increase in FN-SkM.Mes-like progenitor cells during therapy (**Figure S6**). Interestingly, the difference was still detectable, though less exaggerated, in orthotopic PDXs generated from those patient samples (SJRHB00026_X1 and _X2; **Figure S6**). Likewise, PDCs generated from the patient (eRMS-8.1, –8.2, and –8.3) showed a similar pattern with a persistent increase in FN-SkM.Mes-like progenitor expression scores (**Figure S6**). Collectively, these findings demonstrate a treatment-induced selection for the FN-SkM.Mes-like progenitor state in FN-RMS.

**Fig. 5.**
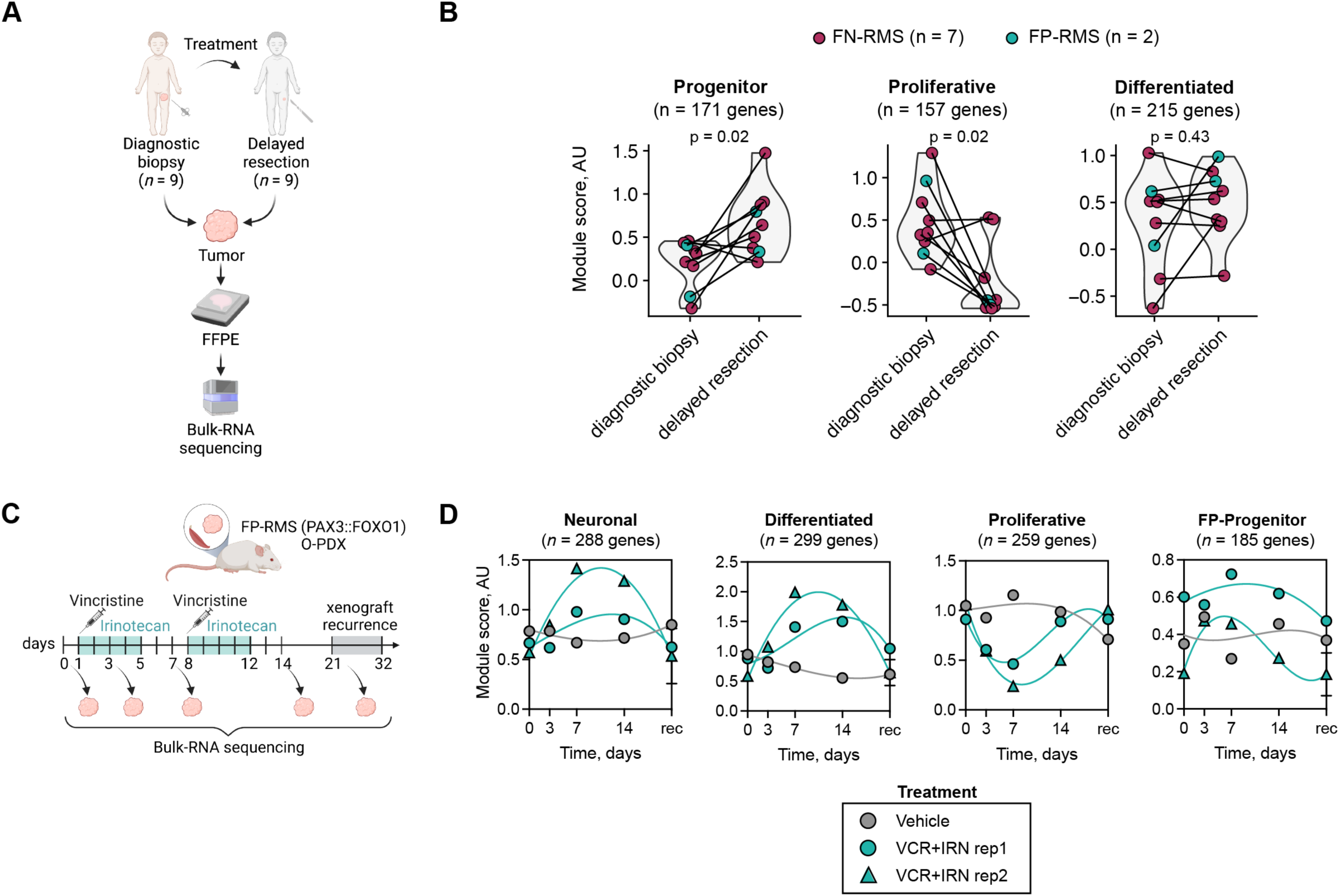
Identification of therapy-resistant RMS metaprograms and cells states in RMS. **A-B**) RNA sequencing analysis of paired diagnostic and on-therapy resection tumors. Experimental design (A). Graphical analysis showing metaprogram scores calculated using the RMS-atlas signature gene sets and shown by tumor subtype (B). Significance noted based on Wilcoxon signed-rank test. **C-D)** RNA-sequencing of tissue obtained from longitudinal biopsies of a FP-RMS orthotopic PDX, SJRHB013759_X14. Experimental design (C). Graphical analysis showing metaprogram scores of bulk RNA sequencing of control treated mouse or those treated with combination of vincristine (VCR) and irinotecan (IRN). Scores were calculated using the signature gene sets identified integrating the FP-RMS PAX3::FOXO1 datasets. Dots were interpolated using a third order polynomial nonlinear fit. AU, arbitrary unit. Marker genes used panel B are from Table S4 while those used in panel D are subtype specific signatures found in Table S5 (PAX3:FOXO1).

Our matched cohort had few FP-RMS patients (*n* = 2; **Figure 5B**), which limited our ability to investigate treatment-induced shifts specifically within FP-RMS. To overcome this limitation, we performed RNA-sequencing of tissue obtained from longitudinal biopsies of a FP-RMS orthotopic PDX, SJRHB013759_X14 [21]. Xenograft-bearing mice were either treated with vehicle or with chemotherapy (vincristine+irinotecan), and sedated needle biopsies were obtained at 5 time points during therapy: day 0 (pre-treatment), day 3, day 7, day 14, and at recurrence (**Figure 5C**). Compared to the vehicle-treated control, tissue obtained from chemotherapy-treated tumors showed upregulation of the differentiated and neuronal signatures at early time points and a return to basal levels at recurrence (**Figure 5D**). Proliferative scores were downregulated at early time points and returned to basal levels at recurrence consistent with the anti-proliferative properties of chemotherapy (**Figure 5D**). In total, these studies identify important new cell states that are retained and expanded after therapy in both FN– and FP-RMS.

## DISCUSSION

Single-cell sequencing technologies have provided unprecedented insight into the intratumoral heterogeneity of a variety of cancers. However, the application of this technology to rare pediatric cancers has been limited by tissue availability, cost, and standardization of bioinformatic analyses. For example, there are ∼350 cases of pediatric rhabdomyosarcoma in the United States annually [2], which limits the ability of any one institute to accrue a sizeable cohort of samples. Here, we combine datasets from groups who published their findings independently to define distinct tumor cell heterogeneity and malignant cell states shared with human muscle cells. This consensus analysis also provides a framework for multi-investigator cooperation that we hope will be an example for future efforts to better understand rare and understudied tumors. Importantly, this work uncovered unexpected new biology related to transitional and therapy resistant cells states and challenged findings across our and others’ previous work, which could only be accomplished with a consensus view of our data and cross-institutional cooperation.

In total, we analyzed 72 samples including 27 FP-RMS, 43 FN-RMS, and 2 MYOD1^L122R^ mutant SS-RMS, encompassing a range of patient tumors, PDXs, PDCs, and commercial cell lines. We identified shared cell states across RMS samples, which we named based on expression similarity shared with human skeletal muscle development including: 1) progenitor cells that express mesenchymal markers and we now name “FN-skeletal muscle mesenchymal-like” (FN-SkM.Mes-like) and “FP-progenitor” cells, 2) differentiated cells that express differentiated muscle-lineage markers, 3) proliferative cells that are enriched for expression of cell cycle genes and largely fail to express progenitor/mesenchymal genes or differentiated muscle genes, and 4) ground state cells that do not show enrichment of any cell state markers. Our new analysis also identified previously underappreciated transitional cell states in both FN– and FP-RMS, suggesting a continuum of states across RMS. Finally, we identify that a subset of FP-RMS harbor small numbers of cells with neural pathway enriched programs and that these same neuronal cells are retained following chemotherapy. These neural cell states were described by Wei et al. previously, but their frequency in FP-RMS had not been reported nor was it known the extent to which PAX7-fusion tumors harbored these cell states [22]. In total, this work provides a standardized nomenclature for RMS cell subpopulations, introduces a transcriptomic muscle-lineage score for assessing cell state, provides cell state signature profiles for harmonizing future studies and analyses of RMS heterogeneity, and confirmed the existence of a tumor-acquired neural cell state in a subset of FP-RMS (**Figure 6**).

**Fig. 6:**
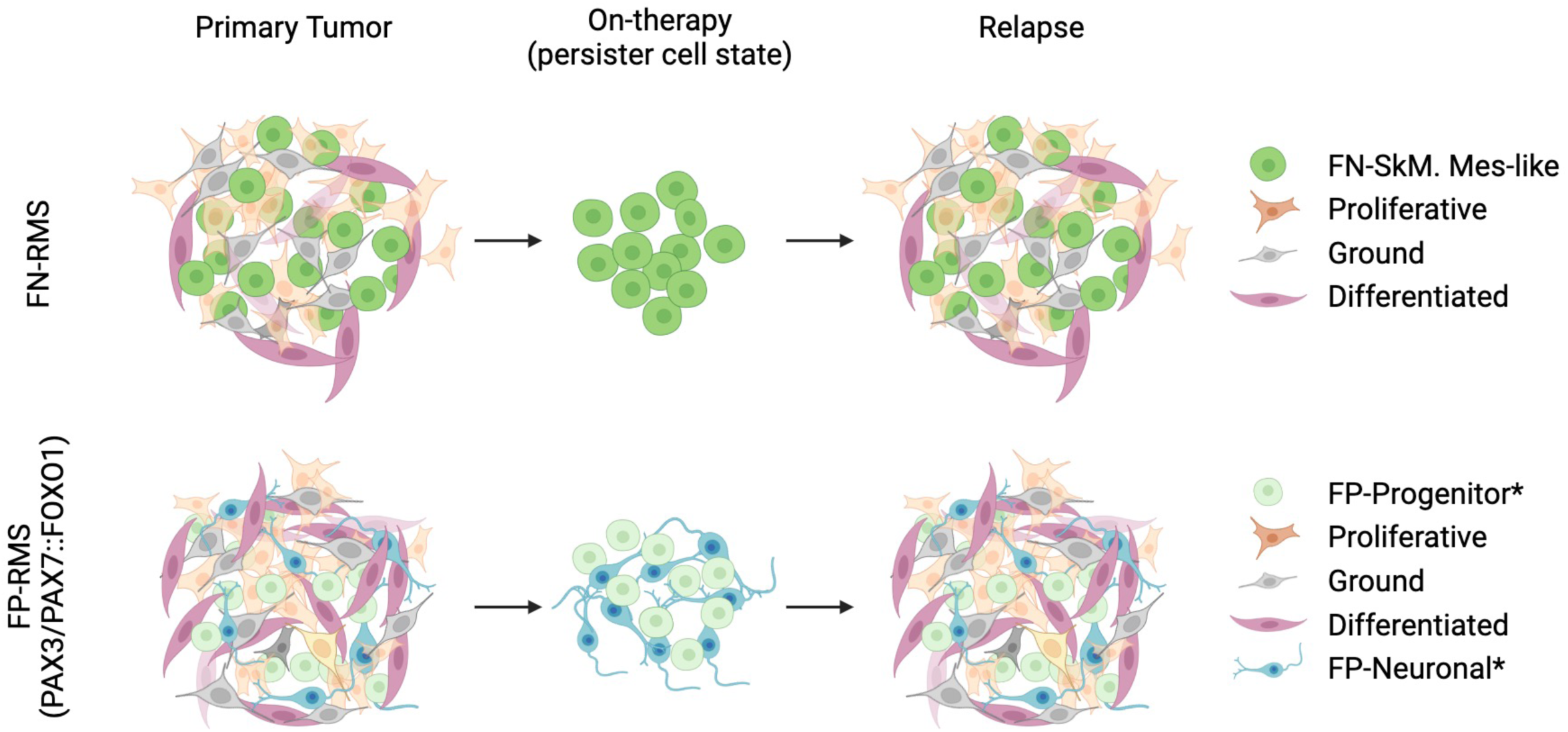
Graphical summary of results. Asterisks denote tumor acquired cells states.

Our findings also contribute to a growing body of literature describing rare cancer cells with the capacity to propagate and re-establish tumors after therapy, which have sometimes been called cancer stem cells or tumor-propagating cells [37–39]. We identified a group of cells expressing a progenitor signature including markers of early muscle progenitors such as *CD44*, *EGFR*, and *THY1* (CD90). Indeed, previous work using flow sorting for cell surface markers such as CD133, CD44 and EGFR have validated the existence of these cells in FN-RMS and demonstrated that they are necessary and sufficient for the propagation of FN-RMS both *in vitro* and when grown in immunocompromised mice [21, 22, 40–44]. Intriguingly, ‘Progenitor’ FN-RMS cells shared gene expression similarity to a newly defined Skeletal muscle mesenchymal cell state that has bipotent capability to make muscle and osteogenic lineage cells and is found only within the second wave of myogenic differentiation in human development [31]. Based on functional studies showing that this RMS cell state can drive tumor growth after stress and has the capacity to make osteogenic lineage cells [22], we have refined our naming of this cell state as “FN-SkM.Mes-like”. Our analysis also suggested that FN-RMS replicate the broad diversity of fetal muscle development cell states and have a shared developmental hierarchy with early developing fetal muscle found after 7-8 weeks post-conception. These findings contrast with Patel *et al* which proposed that FN-RMS recapitulate an earlier mesodermal specification program in mice [21]. This difference is likely attributable to interspecies variation in myogenesis, especially since bipotent SkM.Mes cells have yet to be identified in mice, or that comparison with human muscle development did not include cell types from the earliest stages of mesodermal specification that begin at 24 days post-conception in humans. Finally, our data suggests that FN-SkM.Mes-like cells are largely quiescent and are likely the therapy persistent cells that re-establish tumors after treatment. Indeed, a similar phenomenon where cells with characteristics of progenitors from the hematopoietic, colon, and brain lineages have been proposed to play roles in leukemia, colorectal cancer, and glioblastoma, respectively [45–47].

Our analysis also uncovered that FP-RMS do not display the same rigid developmental hierarchies as found in normal development and may contain different therapy persister cell states. For example, FP-RMS tumors have fewer overall proportions of progenitor cells with some tumors seemingly lacking this cell state completely. Moreover, although the FP-progenitor cells do expresses mesenchymal markers, they are transcriptionally distinct from the SkM.Mesen cells found in fetal muscle development and the FN-SkM.Mes-like state discovered here. Indeed, DeMartino et al., also independently identified key differences in mesenchymal-pathway enriched cell states in comparing scRNA sequencing expression of FP– and FN-RMS [24]. In addition, a subset of FP-RMS have tumor cells that have neural-pathway activation that are not found in human muscle development and yet are enriched after chemotherapy. The existence of this FP-RMS cell state is supported by immunohistochemical studies of 42 FP-RMS tumors that identified a subset of FP-RMS express neuroendocrine marker genes including chromogranin, CD56, and synaptophysin [48]. The frequency by which these tumor-acquired cell states are found in FP-RMS and defining their possible role in driving therapy resistance will clearly be the focus of future work for the field. Our work also suggests that FP-RMS may not follow the rigid-stem cell hierarchies found in development. In fact, limiting dilution cell transplantation assays using engraftment into immune deficient mice have shown that FP-RMS have a high frequency for tumor initiation [49], raising the possibility that most if not all FP-RMS cells can acquire the ability to propagate tumors *in vivo*. Such a model would be consistent with reports where drug-induced neuroendocrine transdifferentiation serves as an escape mechanism for melanoma and castration-resistant prostate cancer [50, 51].

In total, our work has defined new cell state heterogeneity in RMS including identifying a continuum of progenitor and muscle differentiation gene expression, two novel FP-RMS cell states that are not shared in muscle development, and subtype specific, therapy-persistent cell states that likely drive tumor regrowth at relapse.

## METHODS

### Animal experiments

Animal experiments and approvals are described in previous publications [21–23].

### Human Subjects

Formalin-fixed tissues from diagnostic and on-treatment RMS tumors were obtained as part of the RMS 13 trial at St. Jude Children’s Research hospital (NCT018766) for analysis performed in Figure 5A-B.

### scRNAseq/snRNA analysis

#### Public datasets

The 10X Genomics scRNAseq/snRNAseq data was collected from previously published datasets [21–23, 25]. All datasets are available at the NCBI Gene Expression Omnibus (GEO) database under the following accession numbers: GEO: GSE218974 (Danielli et al.; *n* = 17 samples), GEO: GSE195709 (Wei et al.; *n* = 18 samples), GEO: GSE174376 (Patel et al.; *n* = 36 samples), GEO: GSE113660 (Cheng et al.; *n* = 1 sample).

#### Data pre-processing

For samples derived from Wei et al. and Danielli et al., filtered Seurat objects were downloaded from GEO repositories. Those objects were generated as previously described [22, 23] using the 10X Genomics Cell Ranger pipeline (version 3.0.1 in Danielli et al; version 3.1.0 in Wei et al.) to map raw sequencing FASTQ files to the human genome reference (hg19 for patient samples, hg38 for primary cultures) or to both the human hg19 and mouse mm10 references (for PDX samples). Low-quality cells, defined as cells with high mitochondrial ratio (>15% in Danielli et al., >20% in Wei et al.), low expressed gene number (<200 in Danielli et al., <1,000 in Wei et al.), high expressed gene number (>8,000), and PDX cells potentially derived from mice (mouse reads ratio >5% in Wei et al.) were already filtered out.

For samples derived from Patel et al., we generated single-cell Seurat objects following the original pipeline [21]. In short, raw sequencing FASTQ files available from GEO were aligned to the human hg19 (for patient samples) or to the combined human hg19 and mouse mm10 references (for PDX samples) using the 10X Genomics Cell Ranger pipeline (version 3.0.0). Low-quality cells, defined as cells with high mitochondrial ratio (>10%), low (<400) or high expressed gene number (>7,000), were filtered out. We further subset each object to keep only malignant tumor cells, defined based on copy-number variation as described in the original publication (ref. Patel).

For the cell line Rh41, we downloaded the filtered gene-cell matrix available on GEO, that was generated as previously described (add ref. Cheng) using the 10X Genomics Cell Ranger pipeline (version 2.0.1) to map raw sequencing FASTQ files to the human hg38 genome reference. Low-quality cells, defined as cells with high mitochondrial ratio (>15%), low (<200) or high expressed gene number (>8,000) were filtered out.

#### Merging of single-cell transcriptome data

To create the RMS atlas, we first subset each sample to typically *n* = 1500 randomly selected cells (**Table S1**), and then merged raw count matrices using Seurat’s *merge* function. This resulted in a total of *n* = 107,523 cells from *n* = 72 RMS samples (**Table S1**).

To create the three subtype-specific RMS atlases [(1): FN-RMS (*n* = 45 samples); (2): PAX3::FOXO1 FP-RMS (*n* = 15 samples); (3): PAX7::FOXO1 FP-RMS (*n* = 11 samples)], we merged subtype-specific raw count matrices using Seurat’s *merge* function.

#### Normalization and data reduction

After merging, we log-normalized the data, selected the top 2,000 variable features for downstream analyses, and scaled the gene expression. We then performed principal component analysis (PCA) and, based on elbow plot, selected the top n = 15 principal components (PCs) to consider for downstream analysis. To visualize the cells, we reduced the dimensionality of the datasets using Uniform Manifold Approximation and Projection (UMAP).

#### Batch correction and clustering

To remove the batch effects from different samples, we integrated the datasets following Seurat’s integration pipeline (https://satijalab.org/seurat/archive/v3.0/integration.html), which is based on the identification of anchor cells between pairs of datasets. We first normalized and selected *n* = 2,000 variable features for downstream integration from each dataset. We then scaled the data and ran PCA on each object. We identified anchors using reciprocal PCA (RPCA), the suggested option for large datasets, and integrated the datasets using the *IntegrateData* function. We then scaled and centered the gene expression, performed PCA. Based on elbow plot, we then selected the number of PCs to retain for downstream analyses. We built a K-nearest neighbor (KNN) graph, used the Louvain algorithm for clustering the cells (resolution of 0.2-0.3), and visualized the cells using UMAP plots. To identify genes that were enriched within each cluster, we used Seurat’s *FindAllMarkers* function filtering for genes with fold-change >log_2_(0.25) in the subtype-specific datasets and >log_2_(0.3) in the integrated dataset, and expressed in at least 25% of cells in the cluster.

#### Annotation of cell clusters

After clustering, we assigned cell states based on the expression of known markers and gene set enrichment analysis [52–54]. Specifically, we used the marker genes of each cluster as input for Enrichr (https://maayanlab.cloud/Enrichr/)[52–54], and looked at the GO Biological Process 2023 enriched terms. To annotate and collapse the clusters that contained similar lineages, we used the expression of known markers and gene set enrichment analysis [52–54]. For example, clusters 6 and 9 of Fig. 1D both expressed high levels of the muscle differentiation markers *MYOG, MYL4, MYH3*, and were therefore collapsed into one category (« Differentiated »); clusters 8 and 1 both expressed high levels of the collagen and extracellular matrix genes *COL3A1*, *COL1A1*, *FN1*, and were therefore collapsed into one category (« Progenitor »).

#### RMS cell scoring for meta-programs

##### Cell-state specific module scoring

To score each cell based on previously identified metaprograms, we selected the gene markers of the original publications as gene inputs (Ref. **Table S4**). We then assigned cell state-specific module scores using the *AddModuleScore* Seurat’s function. This function works by taking an input set of genes and comparing their average relative expression to that of a control set of *n* = 100 genes randomly sampled [55].

To calculate the consensus *progenitor*, *proliferative*, and *differentiated* marker gene set, we selected cell state markers that were enriched in at least two original publications (Ref. **Table S4**), or in one of the original publications and in the integrated RMS atlas clusters. We then assigned cell state-specific module scores using the *AddModuleScore* Seurat’s function.

We defined the *muscle lineage score* by subtracting the *progenitor* score from the *differentiated* score. Unless otherwise specified, cells were scored using the new consensus *progenitor*, *proliferative*, and *differentiated* markers. The datasets were scaled using the *ScaleData* Seurat’s function to center the expression values.

##### Cell-cycle scoring

After integration, we assigned cell cycle scores using Seurat’s *CellCycleScoring* function, which relies on gene signatures that have been previously shown to characterize S and G2/M cell cycle phases [55]. We distinguished high cycling (S-scores or G2/M scores > 0) from low cycling cells (S-scores < 0 and G2/M scores < 0) based on S and G2/M scores.

#### Comparison of RMS tumors with single-cell reference data from human development

To infer comparisons between RMS tumors and human skeletal muscle development, we re-analyzed a scRNAseq dataset of human skeletal muscle development (GEO: GSE147457) [31]. We downloaded gene expression matrices and their corresponding metadata information for the myogenic subsets derived from embryonic development (1), fetal development (2), juvenile (3) and adult (4) directly from the authors (http://cells.ucsc.edu/?ds=skeletal-muscle). After merging the raw count matrices of the individual datasets, we log-normalized the data, selected the top 2,000 variable features for downstream analyses, and scaled the gene expression. We then performed PCA and, based on elbow plot, selected the top *n* = 10 PCs for downstream analysis. To visualize the cells, we reduced the dimensionality of the datasets using Uniform Manifold Approximation and Projection (UMAP).

To recognize the cell types and developmental time points at which RMS tumors might arise, we used SingleR [36], a computational framework that takes a dataset with known labels as an input and that transfers them onto a test dataset based on similarity to the reference. Specifically, we projected signatures from the human development dataset [31] onto our combined FN-RMS, PAX3::FOXO1 FP-RMS and PAX7::FOXO1 FP-RMS single cell objects.

### Bulk RNA-seq

#### Sample collection

Flash-frozen orthotopic PDX biopsy tissue generated from Patel, et al.[21] underwent RNA isolation using Trizol (Invitrogen) extraction, as per manufacturer instructions. Samples were manually homogenized in 800 μl Trizol within microcentrifuge tubes using a disposable plastic pestle (Fisher Scientific). An additional 200 μl of chloroform was added and mixed via inverting for 3 min at room temperature. Samples were then centrifuged at 12,000 x *g* for 15 minutes at 4 °C. The aqueous layer was transferred to a new microcentrifuge tube, and RNA was precipitated by the addition of an equal volume (approximately 500 μl) of isopropanol and 1 μl glycogen (Thermo Scientific). Samples were incubated at room temperature for 10 minutes, followed by centrifugation at 12,000 x *g* for 15 minutes at 4 °C. Pellets were washed twice with 75% ethanol and resuspended in nuclease-free water. RNA quality was estimated using RNA ScreenTape on a TapeStation automated electrophoresis instrument (Agilent). Sequencing libraries were generated using the TruSeq Total Stranded RNA Library kit (Illumina) using 250-1000 ng of input RNA. Libraries underwent 100 nucleotide paired end sequencing on a NovaSeq 6000 (Illumina).

For extracting RNA from formalin-fixed paraffin-embedded (FFPE) tissue of patient tumors, 5 μm tissue scrolls were processed using the Maxwell RSC RNA FFPE instrument (Promega). RNA concentration and quality was determined using a TapeStation automated electrophoresis instrument (Agilent). Samples with a DV_200_ score (calculated as a percentage of nucleic acid fragments > 200 nucleotides) above 20% were used for downstream library generation using the SMARTer Stranded Total RNA – Pico RNA-seq kit v2 (Takara). Libraries were sequenced using 100 nucleotide paired-end sequencing on a NovaSeq 6000 (Illumina).

#### Data pre-processing

Following sequencing, all RNA-seq libraries were processed using an automated computational pipeline. Briefly, sequenced reads were trimmed and underwent quality control using FastQC, followed by aligning and counting using STAR [56]. To generate additional alignment metrics, the STAR-aligned BAM file were analyzed using the ’CollectRNASeqMetrics’ command from the Picard pipeline. Duplicate reads were removed using the GATK ‘MarkDupliates’ command, and then RSEM was used to generate tpm count matrices using the ‘rsem-calculate-expression’ command.

#### Bulk-RNAseq scoring

To score FFPE tissues and orthotopic PDX biopsies for the *progenitor*, *proliferative* or *differentiated* scores, we first created a Seurat object using the already TPM-normalized read count matrix, and log-normalized the count matrix expression values+1. We then scored individual samples for the cell state-specific module scores using the *AddModuleScore* Seurat’s function. We plotted the scores after scaling and centering the expression values using the *ScaleData* Seurat’s function.

## DATA AVAILABILITY

Published single-cell/nucleus RNA-sequencing data were obtained from the Gene Expression Omnibus (GEO): Danielli, et al. (GSE218974); Patel, et al. (GSE174376); Wei, et al (GSE195709); Cheng, et al. (GSE113660). RNA-sequencing data generated from matched patient samples before and during therapy as well as biopsied FP-RMS orthotopic PDXs will be available upon publication (GSE240287 and GSE240308, respectively).

## ACKNOWLEDGMENTS

This work was funded by the Sarcoma Foundation of America (2022 SFA 13-22, B.W.S. and S.G.D.), the Hyundai Hope on Wheels Foundation (A.G.P.), the Damon Runyon Cancer Foundation (#DRSG-33P-20, A.G.P.). the Alex’s Lemonade Stand Foundation (M.A.D. and A.G.P.), CureSearch (D.M.L), American Lebanese Syrian Associated Charities, the Friends of TJ and Summer’s Way Foundation (Y.W), MGH ECOR Medical Discovery Award (Y.W), the Rally Foundation (D.M.L.), and the NCI (K99CA278696 (Y.W), R01CA276116 (D.M.L), R01CA269213 (D.M.L), U54CA231630 (D.M.L.)). We thank the St. Jude Clinical Biomarkers Laboratory for assistance with RNA extraction of formalin-fixed paraffin embedded tissues, the St. Jude Hartwell Center for Biotechnology for sequencing support, and the St. Jude Center for Applied Bioinformatics for support with RNA-sequencing analysis. Some figures were created with BioRender.com.

## AUTHOR CONTRIBUTIONS

D.M.L., A.G.P., M.W., and B.W.S supervised the study design and writing. S.G.D. performed the analysis and generated figures. S.G.D. and Y. W. coordinated the collaboration, provided feedback on ideas and figures, and wrote the manuscript. M.A.D. and E.S. provided samples for the analysis of treatment-induced expression signatures.

## REFERENCES

1. Kashi, V.P., M.E. Hatley, and R.L. Galindo, Probing for a deeper understanding of rhabdomyosarcoma: insights from complementary model systems. Nat Rev Cancer, 2015. 15(7): p. 426–39.

2. Skapek, S.X., et al., Rhabdomyosarcoma. Nat Rev Dis Primers, 2019. 5(1): p. 1.

3. Dasgupta, R., J. Fuchs, and D. Rodeberg, Rhabdomyosarcoma. Semin Pediatr Surg, 2016. 25(5): p. 276–283.

4. Davicioni, E., et al., Identification of a PAX-FKHR gene expression signature that defines molecular classes and determines the prognosis of alveolar rhabdomyosarcomas. Cancer Res, 2006. 66(14): p. 6936–46.

5. Davicioni, E., et al., Molecular classification of rhabdomyosarcoma--genotypic and phenotypic determinants of diagnosis: a report from the Children’s Oncology Group. Am J Pathol, 2009. 174(2): p. 550–64.

6. Fletcher, C.D.M., et al., WHO classification of tumours of soft tissue and bone. 2013.

7. Rekhi, B., et al., MYOD1 (L122R) mutations are associated with spindle cell and sclerosing rhabdomyosarcomas with aggressive clinical outcomes. Mod Pathol, 2016. 29(12): p. 1532–1540.

8. Shern, J.F., et al., Genomic Classification and Clinical Outcome in Rhabdomyosarcoma: A Report From an International Consortium. J Clin Oncol, 2021. 39(26): p. 2859–2871.

9. Kohsaka, S., et al., A recurrent neomorphic mutation in MYOD1 defines a clinically aggressive subset of embryonal rhabdomyosarcoma associated with PI3K-AKT pathway mutations. Nat Genet, 2014. 46(6): p. 595–600.

10. Agaram, N.P., et al., MYOD1-mutant spindle cell and sclerosing rhabdomyosarcoma: an aggressive subtype irrespective of age. A reappraisal for molecular classification and risk stratification. Mod Pathol, 2019. 32(1): p. 27–36.

11. Alaggio, R., et al., A Molecular Study of Pediatric Spindle and Sclerosing Rhabdomyosarcoma: Identification of Novel and Recurrent VGLL2-related Fusions in Infantile Cases. Am J Surg Pathol, 2016. 40(2): p. 224–35.

12. Butel, T., et al., Integrative clinical and biopathology analyses to understand the clinical heterogeneity of infantile rhabdomyosarcoma: A report from the French MMT committee. Cancer Med, 2020. 9(8): p. 2698–2709.

13. Mosquera, J.M., et al., Recurrent NCOA2 gene rearrangements in congenital/infantile spindle cell rhabdomyosarcoma. Genes Chromosomes Cancer, 2013. 52(6): p. 538–50.

14. Mascarenhas, L., et al., Randomized phase II window trial of two schedules of irinotecan with vincristine in patients with first relapse or progression of rhabdomyosarcoma: a report from the Children’s Oncology Group. J Clin Oncol, 2010. 28(30): p. 4658–63.

15. Pappo, A.S., et al., Survival after relapse in children and adolescents with rhabdomyosarcoma: A report from the Intergroup Rhabdomyosarcoma Study Group. J Clin Oncol, 1999. 17(11): p. 3487–93.

16. Smith, L.M., et al., Which patients with microscopic disease and rhabdomyosarcoma experience relapse after therapy? A report from the soft tissue sarcoma committee of the children’s oncology group. J Clin Oncol, 2001. 19(20): p. 4058–64.

17. Chisholm, J.C., et al., Prognostic factors after relapse in nonmetastatic rhabdomyosarcoma: a nomogram to better define patients who can be salvaged with further therapy. J Clin Oncol, 2011. 29(10): p. 1319–25.

18. Grobner, S.N., et al., The landscape of genomic alterations across childhood cancers. Nature, 2018. 555(7696): p. 321–327.

19. Chen, L., et al., Clonality and evolutionary history of rhabdomyosarcoma. PLoS Genet, 2015. 11(3): p. e1005075.

20. Chen, X., et al., Targeting oxidative stress in embryonal rhabdomyosarcoma. Cancer Cell, 2013. 24(6): p. 710–24.

21. Patel, A.G., et al., The Myogenesis Program Drives Clonal Selection and Drug Resistance in Rhabdomyosarcoma. bioRxiv, 2021: p. 2021.06.16.448386.

22. Wei, Y., et al., Single-cell analysis and functional characterization uncover the stem cell hierarchies and developmental origins of rhabdomyosarcoma. Nat Cancer, 2022. 3(8): p. 961–975.

23. Danielli, S.G., et al., Single-cell profiling of alveolar rhabdomyosarcoma reveals RAS pathway inhibitors as cell-fate hijackers with therapeutic relevance. Sci Adv, 2023. 9(6): p. eade9238.

24. DeMartino, J., et al., Single-cell transcriptomics reveals immune suppression and cell states predictive of patient outcomes in rhabdomyosarcoma. Nat Commun, 2023. 14(1): p. 3074.

25. Cheng, C., et al., Latent cellular analysis robustly reveals subtle diversity in large-scale single-cell RNA-seq data. Nucleic Acids Res, 2019. 47(22): p. e143.

26. Stewart, E., et al., Orthotopic patient-derived xenografts of paediatric solid tumours. Nature, 2017. 549(7670): p. 96–100.

27. Neftel, C., et al., An Integrative Model of Cellular States, Plasticity, and Genetics for Glioblastoma. Cell, 2019. 178(4): p. 835–849 e21.

28. Wu, S.Z., et al., A single-cell and spatially resolved atlas of human breast cancers. Nat Genet, 2021. 53(9): p. 1334–1347.

29. Izar, B., et al., A single-cell landscape of high-grade serous ovarian cancer. Nat Med, 2020. 26(8): p. 1271–1279.

30. Hao, Y., et al., Integrated analysis of multimodal single-cell data. Cell, 2021. 184(13): p. 3573–3587 e29.

31. Xi, H., et al., A Human Skeletal Muscle Atlas Identifies the Trajectories of Stem and Progenitor Cells across Development and from Human Pluripotent Stem Cells. Cell Stem Cell, 2020. 27(1): p. 181–185.

32. Castiglioni, A., et al., Isolation of progenitors that exhibit myogenic/osteogenic bipotency in vitro by fluorescence-activated cell sorting from human fetal muscle. Stem Cell Reports, 2014. 2(1): p. 92–106.

33. Chal, J. and O. Pourquie, Making muscle: skeletal myogenesis in vivo and in vitro. Development, 2017. 144(12): p. 2104–2122.

34. Bentzinger, C.F., Y.X. Wang, and M.A. Rudnicki, Building muscle: molecular regulation of myogenesis. Cold Spring Harb Perspect Biol, 2012. 4(2).

35. Shi, X. and D.J. Garry, Muscle stem cells in development, regeneration, and disease. Genes Dev, 2006. 20(13): p. 1692–708.

36. Aran, D., et al., Reference-based analysis of lung single-cell sequencing reveals a transitional profibrotic macrophage. Nat Immunol, 2019. 20(2): p. 163–172.

37. Genadry, K.C., et al., Soft Tissue Sarcoma Cancer Stem Cells: An Overview. Front Oncol, 2018. 8: p. 475.

38. Dela Cruz, F.S., Cancer stem cells in pediatric sarcomas. Front Oncol, 2013. 3: p. 168.

39. Hettmer, S. and A.J. Wagers, Muscling in: Uncovering the origins of rhabdomyosarcoma. Nat Med, 2010. 16(2): p. 171–3.

40. Walter, D., et al., CD133 positive embryonal rhabdomyosarcoma stem-like cell population is enriched in rhabdospheres. PLoS One, 2011. 6(5): p. e19506.

41. Radzikowska, J., et al., Cancer Stem Cell Markers in Rhabdomyosarcoma in Children. Diagnostics (Basel), 2022. 12(8).

42. Linardic, C.M., et al., Genetic modeling of human rhabdomyosarcoma. Cancer Res, 2005. 65(11): p. 4490–5.

43. Ignatius, M.S., et al., In vivo imaging of tumor-propagating cells, regional tumor heterogeneity, and dynamic cell movements in embryonal rhabdomyosarcoma. Cancer Cell, 2012. 21(5): p. 680–693.

44. Blum, J.M., et al., Distinct and overlapping sarcoma subtypes initiated from muscle stem and progenitor cells. Cell Rep, 2013. 5(4): p. 933–40.

45. Singh, S.K., et al., Identification of human brain tumour initiating cells. Nature, 2004. 432(7015): p. 396–401.

46. Ricci-Vitiani, L., et al., Identification and expansion of human colon-cancer-initiating cells. Nature, 2007. 445(7123): p. 111–5.

47. Lapidot, T., et al., A cell initiating human acute myeloid leukaemia after transplantation into SCID mice. Nature, 1994. 367(6464): p. 645–8.

48. Bahrami, A., et al., Aberrant expression of epithelial and neuroendocrine markers in alveolar rhabdomyosarcoma: a potentially serious diagnostic pitfall. Mod Pathol, 2008. 21(7): p. 795–806.

49. Generali, M., et al., High Frequency of Tumor Propagating Cells in Fusion-Positive Rhabdomyosarcoma. Genes (Basel), 2021. 12(9).

50. Zou, M., et al., Transdifferentiation as a Mechanism of Treatment Resistance in a Mouse Model of Castration-Resistant Prostate Cancer. Cancer Discov, 2017. 7(7): p. 736–749.

51. Rambow, F., et al., Toward Minimal Residual Disease-Directed Therapy in Melanoma. Cell, 2018. 174(4): p. 843–855 e19.

52. Kuleshov, M.V., et al., Enrichr: a comprehensive gene set enrichment analysis web server 2016 update. Nucleic Acids Res, 2016. 44(W1): p. W90–7.

53. Chen, E.Y., et al., Enrichr: interactive and collaborative HTML5 gene list enrichment analysis tool. BMC Bioinformatics, 2013. 14: p. 128.

54. Xie, Z., et al., Gene Set Knowledge Discovery with Enrichr. Curr Protoc, 2021. 1(3): p. e90.

55. Tirosh, I., et al., Dissecting the multicellular ecosystem of metastatic melanoma by single-cell RNA-seq. Science, 2016. 352(6282): p. 189–96.

56. Dobin, A., et al., STAR: ultrafast universal RNA-seq aligner. Bioinformatics, 2013. 29(1): p. 15–21.

