## Supplementary figures and legends for "Single cell transcriptomic profiling identifies tumor-acquired and therapy-resistant cell states in pediatric rhabdomyosarcoma"

* denotes equal authorship

\# corresponding authors

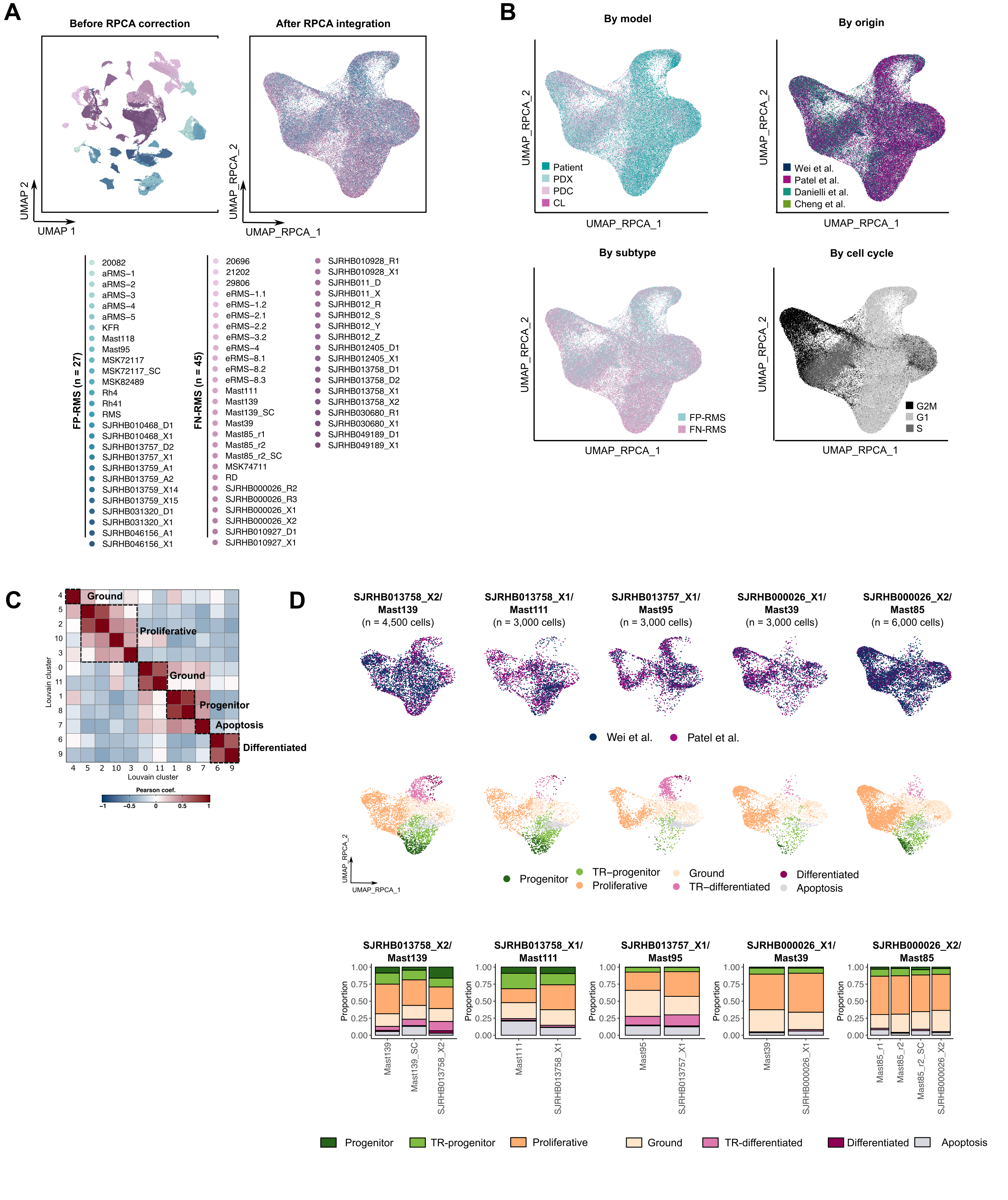
**Fig. S1. Integrated analysis of single cell and single nuclei sequencing of human RMS. A)** UMAP rendering of RMS cells before (left) and after RPCA integration (right). Cells are colored based on sample as noted in the legend below the PCA plot. “_SC”, single-cell derived PDX sample, “_D”, diagnostic, “_R”, recurrence, “_A”, autopsy, “_X”. xenograft. **B)** UMAP rendering of RMS cells colored by model, publication of origin, RMS subtype, or cell cycle. **C)** Heatmap showing correlation matrix similarities of Louvain clusters identified by integrated analysis of all RMS samples together. Clusters were combined based on gene expression similarity and cell states are noted by dashed lines. **D)** Comparison of relative cell state constitution across five PDX models sequenced from different publications. UMAP plots (top) and bar plots showing cell state composition (bottom).

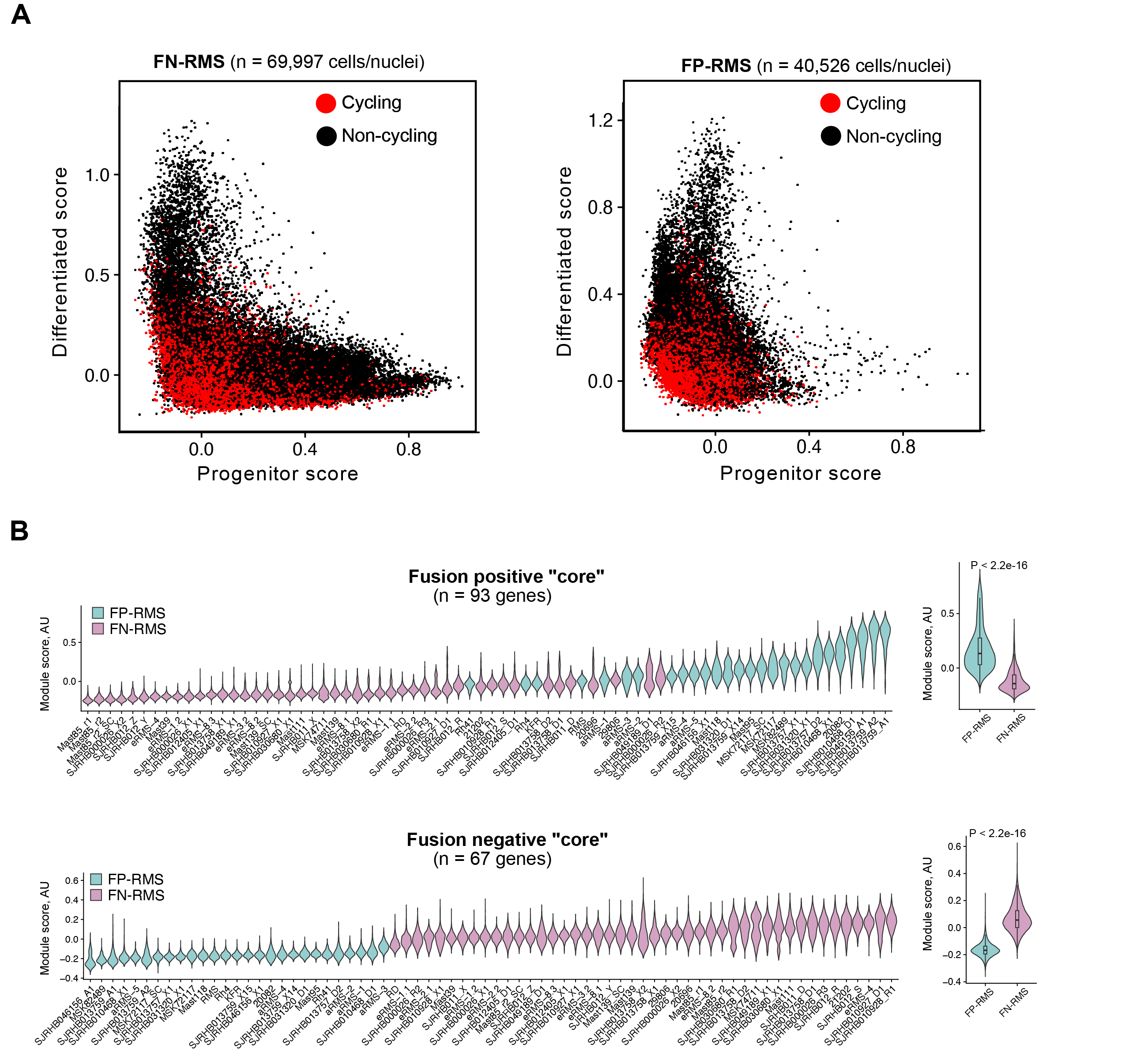

**Fig. S2. Gene expression across single cells from FN- and FP- RMS. A)** Scatter plot comparing individual cell expression of progenitor and differentiated metaprograms across FN-RMS (left) or FP-RMS samples (right). Cells/nuclei are colored by cycling status. **B)** Violin plots showing expression of core-signature gene profiles that distinguish the two major RMS subtypes as defined by Wei *et al.* 2022. Left: Analysis of each tumor model. Right: Summary analysis after combining all samples together. AU, arbitrary unit. Statistical analysis used Student’s T-test comparison with p-values noted.

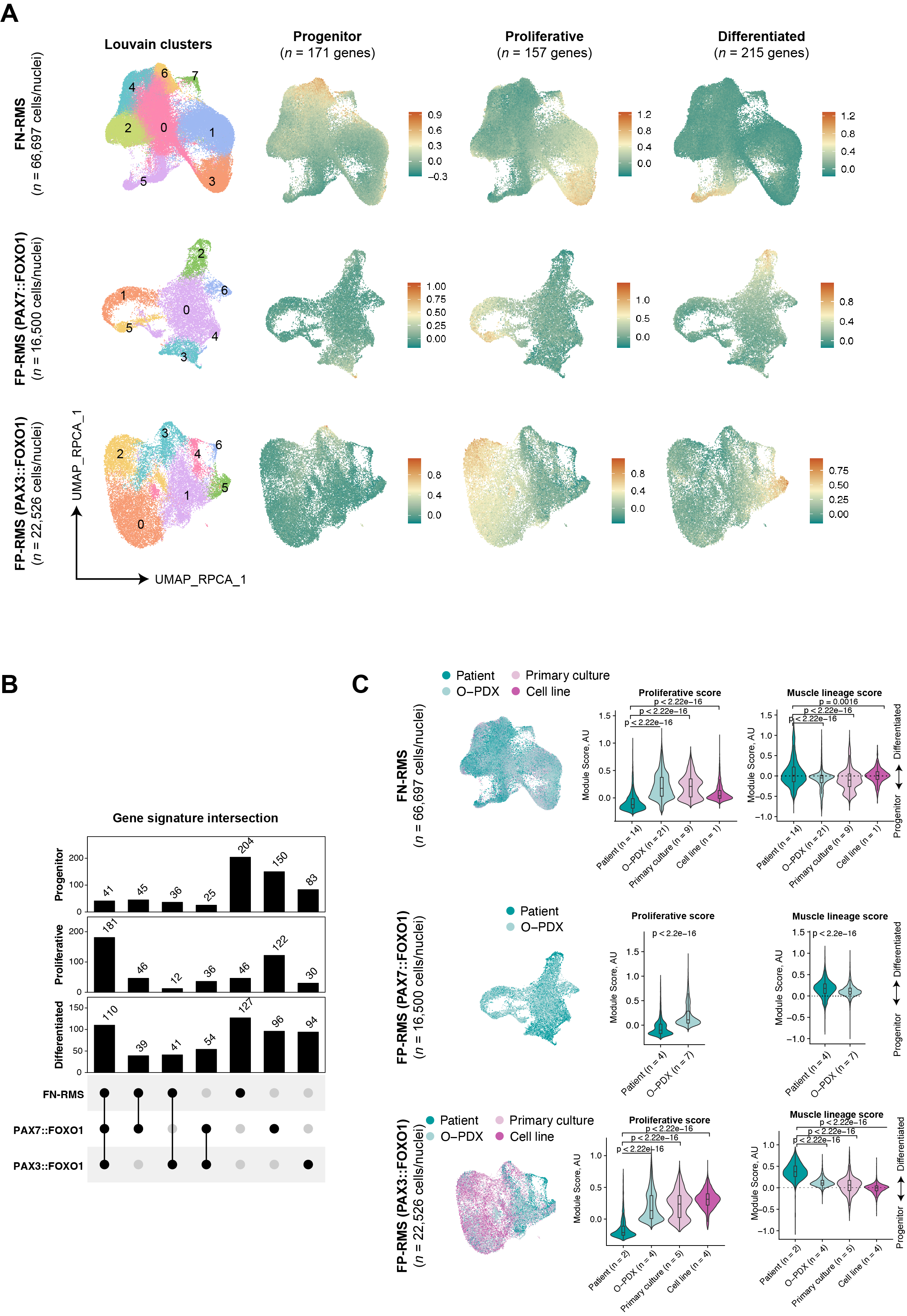

**Fig. S3. Tripartite muscle intra-tumoral heterogeneity is largely retained across patient and preclinical RMS models. A**) UMAP renderings of FN-RMS, PAX7::FOXO1 and PAX3::FOXO1 FP-RMS. Cells/nuclei were integrated independently and colored based on Louvian clusters (left) or by the expression of cell state metaprograms defined in this publication (right). **B**) Upset plot of the marker genes for the progenitor, proliferative and differentiated signatures derived from the subtype-specific analysis. The progenitor signature shows little overlap among the three RMS subtypes (only 41 genes), whereas the proliferative and differentiated signatures share a higher fraction of overlapping genes across models. **C)** Analysis of tripartite muscle cell states in patient tissue, orthotopic xenografts, primary cultures, or established cell lines. UMAP plots comparing RMS subtype (left) and Violin plot quantitation (right). Adjusted *p*-values calculated by one-way ANOVA (FN-RMS and PAX3::FOXO1 FP-RMS) or by Student t-test (PAX7::FOXO1 FP-RMS).

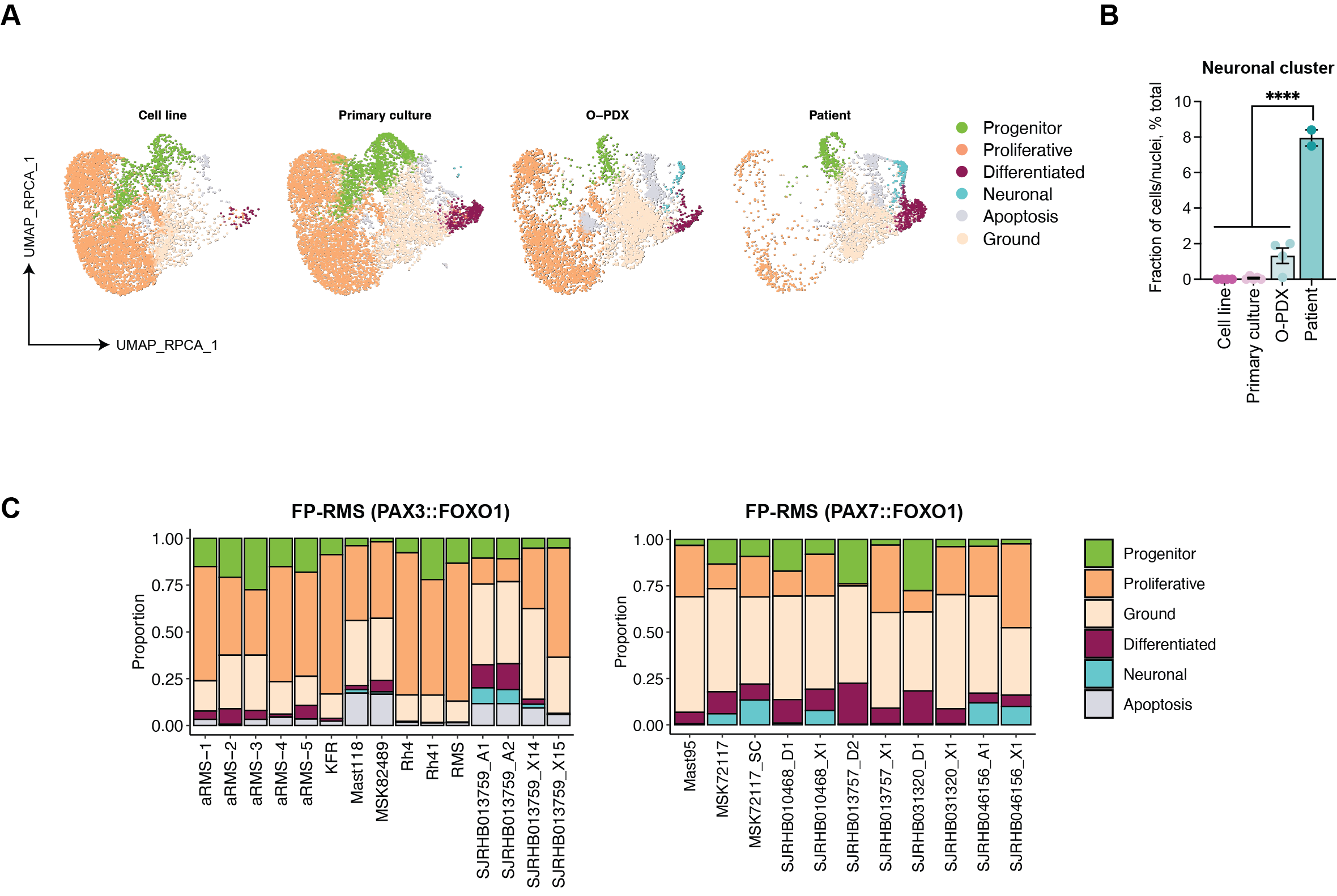
**Fig. S4. Neural cell states are not retained in FP-RMS cell line and primary cultures. A)** UMAP plots of integrated PAX3::FOXO1 FP-RMS samples colored by cell states and analyzed across tumor models **B)** Quantification of the neuronal cluster fraction across PAX3::FOXO1 FP-RMS samples. Ordinary one-way analysis of variance (ANOVA) with Dunnett’s multiple comparison correction. ****P ≤ 0.0001).

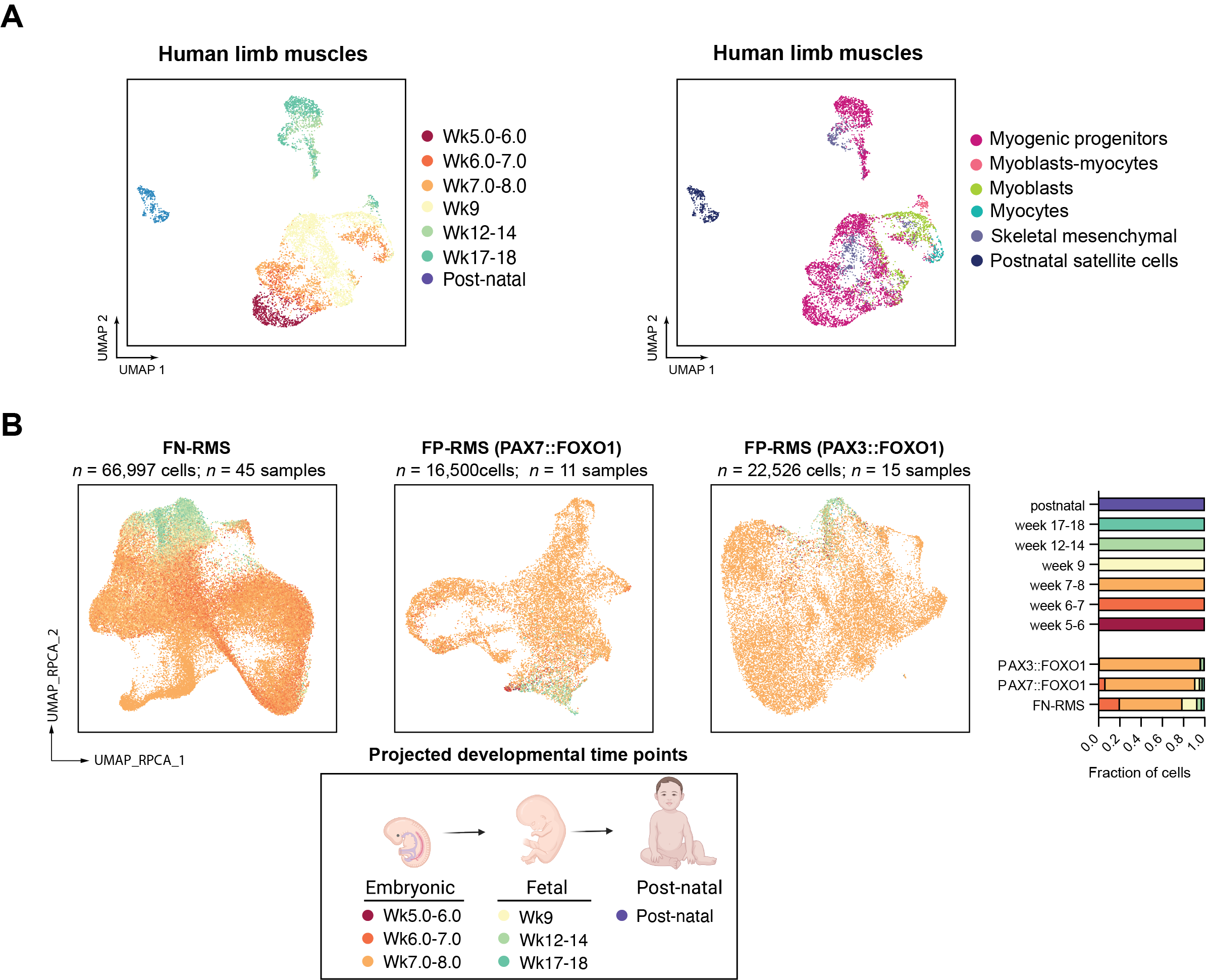

**Fig. S5: Mapping RMS cell similarity based on developmental age. A)** UMAP plots for developmental muscle atlas used in this analysis (Xi *et al.* 2020*. Cell Stem Cell*). Cells are colored by developmental time point (left) or cell type (right). Post-natal samples include tissues obtained from skeletal muscle of 7 to 42 year olds. B**)** UMAP plots of RMS fusion subtype and colored by shared developmental time point similarity to muscle using SingleR (left) and quantification (right). Wk, week.

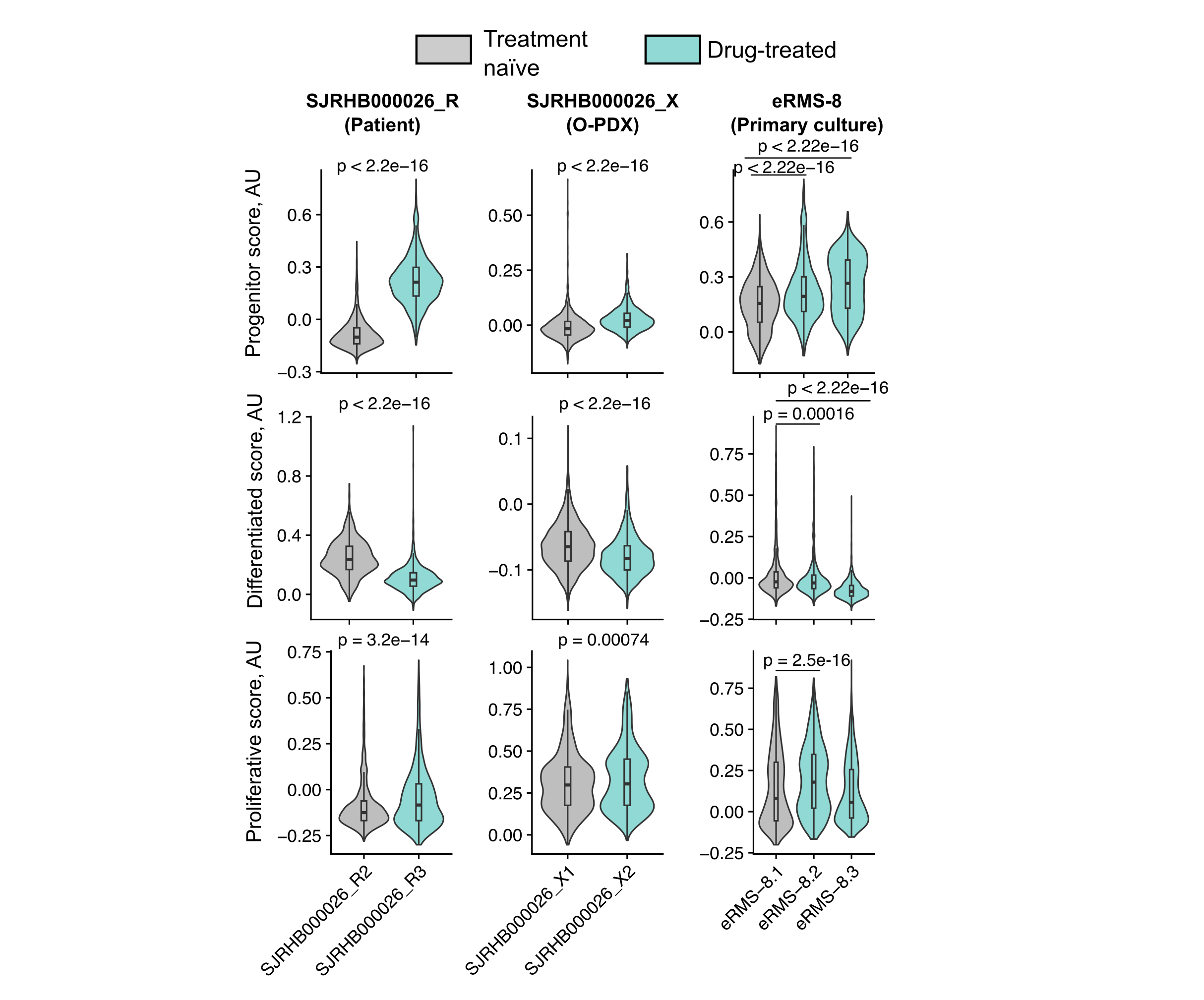
**Fig. S6. FN-RMS progenitor cells are increased following chemotherapy treatment.** Metaprogram scores assigned across cells/nuclei derived from paired primary patient, O-PDX and primary cultures established from tumors before (treatment naïve) and during treatment (drug-treated, vincristine+irinotecan)). *P*-values were calculated by Student’s t test (SJRHB000026_R and SJRHB000026_X) or one-way ANOVA (eRMS-8).

**SUPPLEMENTARY TABLES**

**Table S1. Clinical properties of the RMS samples used for this analysis.**

**Table S2. Marker genes differentially expressed among the identified Seurat clusters in the integrated RMS atlas (FC>0.3).**

**Table S3. Gene enrichment analysis of the marker genes differentially expressed among Seurat clusters in the integrated RMS atlas (FC>0.3).**

**Table S4. Marker genes for the progenitor, proliferating and differentiated cell states from the original publications and this new analysis.**

**Table S5. Cluster markers identified across the FN-RMS, FP-RMS (PAX3::FOXO1) and FP-RMS (PAX7::FOXO1) RMS subtype-specific datasets.** Marker genes differentially expressed among the identified Seurat clusters (FC>0.25).

**Table S6. Gene enrichment analysis of the marker genes differentially expressed among clusters in subtype specific analysis of FN-RMS, FP-RMS (PAX3::FOXO1) and FP-RMS (PAX7::FOXO1) RMS.** (FC>0.25)

**Table S7. Frequency of cells across the clusters identified in the FN-RMS, FP-RMS (PAX3::FOXO1) and FP-RMS (PAX7::FOXO1) RMS subtype-specific datasets.** Not detected (ND).

**Table S8. Number and fraction of RMS cells mapping to the developmental age and cell types found in normal human muscle.** Muscle data set from Xi et al., Cell Stem Cell (2020).
